# Catecholaminergic modulation of the avoidance of cognitive control

**DOI:** 10.1101/191015

**Authors:** Monja I. Froböse, Jennifer C. Swart, Jennifer L. Cook, Dirk E.M. Geurts, Hanneke E.M. den Ouden, Roshan Cools

## Abstract

The catecholamines have long been associated with cognitive control and value-based decision-making. More recently, we proposed that the catecholamines might modulate value-based decision-making about whether or not to engage in cognitive control. We test this hypothesis by assessing effects of a catecholamine challenge in a large sample of young, healthy adults (n = 100) on the avoidance of a cognitively demanding control process: task switching. Prolonging catecholamine transmission by blocking reuptake with methylphenidate altered the avoidance, but not the execution of cognitive control. Crucially, these effects could be isolated by taking into account individual differences in trait impulsivity, so that participants with higher trait impulsivity became more avoidant of cognitive control, despite faster task performance. One implication of these findings is that performance-enhancing effects of methylphenidate may be accompanied by an undermining effect on the willingness to exert cognitive control. Taken together, these findings integrate hitherto segregated literatures on catecholamines’ roles in value-based learning/choice and cognitive control.

## INTRODUCTION

Catecholamine neurotransmitters (dopamine and noradrenaline) have long been implicated in key aspects of goal-directed behaviour, including on the one hand cognitive control (Arnsten, 1998; Brozoski, Brown, Rosvold, & Goldman, 1979; Cools, D’Esposito, 2011; Cools, Clark, & Robbins, 2004; Goldman-Rakic, 1997) and on the other hand value-based learning, motivation and choice (Collins & Frank, 2014; Niv, Daw, Joel, & Dayan, 2007; Robbins & Everitt, 1996; Salamone, Correa, Mingote, & Weber, 2005; Schultz, 2017). Recently, catecholamines have been proposed to also mediate their integration: value-based learning and choice about whether or not to recruit cognitive control (Cools, 2016; Westbrook & Braver, 2016). This idea implies that catecholaminergic drugs, such as methylphenidate (MPH), alter not just the ability to execute cognitive control, but also the willingness to exert or conversely, the desire to avoid, cognitive control. Here, we test this hypothesis by assessing the effects of a catecholamine challenge on the avoidance of cognitive control.

### Catecholaminergic modulation of cognitive control

Cognitive control refers to the ability to flexibly adjust our behaviour to changing internal and external demands in order to attain (long-term) goals (Egner, 2017; Fuster, 1989; Monsell, 2003). Disorders accompanied by cognitive control deficits, such as attention deficit/ hyperactivity disorder (ADHD), Parkinson’s disease and schizophrenia, are commonly treated with drugs that alter catecholamine transmission (Arnsten, 1998; Dagher & Robbins, 2009; Frankle & Laruelle, 2002; Prince, 2008). In ADHD, for example, MPH is usually the first-line medication and is generally found to remedy cognitive deficits (Coghill et al., 2013; Faraone & Buitelaar, 2010; Leonard, Mccartan, White, & King, 2004), such as impairments in task switching (Cepeda, Cepeda, & Kramer, 2000; Kramer, Cepeda, & Cepeda, 2001), response inhibition (Aron, Dowson, Sahakian, & Robbins, 2003), and working memory (Mehta, Goodyer, & Sahakian, 2004). In addition, psychostimulants, such as MPH have been shown to enhance cognitive function in healthy volunteers (Linssen, Sambeth, Vuurman, & Riedel, 2014), consistent with their use by students and academics to boost functioning in periods of high cognitive demand (Maher, 2008). Acute administration of a single dose of psychostimulants to healthy volunteers has indeed been shown to improve task switching (Samanez-Larkin & Buckholtz, 2013), extradimensional set-shifting (Rogers et al., 1999), spatial working memory (Elliott et al., 1997), response inhibition (Spronk, Bruijn, Wel, Ramaekers, & Verkes, 2013), distractor-resistant working memory (Fallon et al., 2016) and selective attention (Ter Huurne et al., 2015). Thus, catecholaminergic drugs can both remedy cognitive control deficits in patients and enhance cognitive control in the healthy population.

However, while drugs that potentiate catecholamine neurotransmission, like MPH, are generally thought to enhance cognitive control, they certainly do not have enhancing effects in all people. Indeed, there is large individual variability in the direction and extent of catecholaminergic drug effects on human cognition (Cools et al., 2004; Samanez-Larkin & Buckholtz, 2013). These individual differences in drug effects are thought to reflect dependency on baseline levels of dopamine (Cools & D’Esposito, 2011) and covary with proxy variables, such as trait impulsivity (associated with dopamine receptor availability and striatal dopamine release; Buckholtz et al., 2010; Kim et al., 2013; Lee et al., 2009; Reeves et al., 2012) and working memory capacity (associated with dopamine synthesis capacity; Cools, Gibbs, Miyakawa, Jagust, & D’Esposito, 2008; Landau et al., 2009). Participants with higher trait impulsivity have been shown to exhibit greater beneficial effects of catecholaminergic drug administration across tasks including attention switching (Cools, Sheridan, Jacobs, & D’Esposito, 2007) and probabilistic reversal learning (Clatworthy et al., 2009). Such impulsivity-dependent effects of catecholaminergic drugs correspond well with the cognitive enhancing effects of MPH in ADHD (Rapoport et al., 1980), with greater MPH-induced changes in dopamine release in more severely affected ADHD patients (Rosa-Neto et al., 2005) and with greater beneficial effects of MPH on impulsive responding in higher impulsive experimental rodents (Caprioli et al., 2015). Thus, we expected that MPH-effects on cognitive control can be isolated by taking into account proxy variables, such as individual trait impulsivity.

### Catecholaminergic modulation of learning and choice about cognitive control

In parallel, a second, so far relatively segregated line of evidence supports a key role for the catecholamines, dopamine in particular, in value-based learning and choice (Collins & Frank, 2014; Cools, Nakamura, & Daw, 2011; Maia & Frank, 2015; Schultz, 2001; Swart et al., 2017; van der Schaaf, Fallon, Ter Huurne, Buitelaar, & Cools, 2013). It is well-established that phasic firing of midbrain dopamine neurons contributes to the encoding of reward prediction errors (Montague, Dayan, & Sejnowski, 1996; Schultz, 1997; Tobler, Fiorillo, & Schultz, 2005), driving reinforcement learning and consequently promoting the selection of actions with higher predicted values. It has been argued that the same principle applies to the selection of cognitive goals, such that dopaminergic reward prediction error signals can contribute to the value-based learning and selection of cognitive goals (Braver & Cohen, 1999; Collins & Frank, 2014; Frank, Loughry, & O’Reilly, 2001; Frank & Badre, 2012; Hazy, Frank, & O’Reilly, 2007).

This evidence concurs with recent expected value accounts of cognitive control (Botvinick & Braver, 2015; Kool, Gershman, & Cushman, 2017; Kurzban, Duckworth, Kable, & Myers, 2013; Shenhav et al., 2017), which propose that the degree (and intensity) of engagement in an upcoming cognitive computation is based on a cost-benefit analysis. In line with this account, it has been shown repeatedly that enhancing motivation, for example by offering reward, affects performance on cognitive control paradigms (Aarts, van Holstein, & Cools, 2011; Botvinick & Braver, 2015; Chib, Shimojo, & O’Doherty, 2014; Chib, De Martino, Shimojo, & O’Doherty, 2012; Manohar et al., 2015; Padmala & Pessoa, 2011). Increasing the value or benefit of a demanding computation, such as task switching, seems to outweigh perceived demand costs.

Evidence is accumulating that cognitive demand indeed carries an intrinsic cost (Botvinick & Braver, 2015; Westbrook & Braver, 2016), a hypothesis that is supported by studies showing that, on average, healthy participants are demand avoidant. They prefer to perform a task with a lower cognitive demand, such as less task switching (Botvinick, 2007; Gold et al., 2015; Kool et al., 2010; McGuire & Botvinick, 2010) or lower working memory load, they choose to forego a higher monetary reward to avoid a more demanding task (Massar, Libedinsky, Weiyan, Huettel, & Chee, 2015; Westbrook, Kester, & Braver, 2013) and expend physical effort in order to reduce cognitive demand (Risko, Medimorec, Chisholm, & Kingstone, 2014).

A role for the catecholamines in biasing meta-learning and -choice about cognitive effort follows also from abundant evidence implicating dopamine in physical effort avoidance. Enhancing dopamine transmission in human and non-human animals increases selection of high effort/high reward trials (Chong et al., 2015; Floresco, Tse, & Ghods-Sharifi, 2008; Le Bouc et al., 2016; Salamone, Yohn, & Lo, 2016), while the opposite is true for reductions in dopamine functioning (e.g. Bardgett, Depenbrock, Downs, & Green, 2009; see Salamone et al., 2016 for review). In these studies, it is evident that dopamine manipulations altered effort-based choice rather than the capacity to exert effort per se because animals were still equally able to execute the physical effortful task of climbing a barrier (Cousins, Atherton, Turner, & Salamone, 1996; Yohn et al., 2015).

There is suggestive empirical evidence that similar mechanisms underlie learning and choice about cognitive demand (Botvinick, Huffstetler, & McGuire, 2009; Kurniawan, Guitart-Masip, Dayan, & Dolan, 2013): Prolonging catecholamine transmission by amphetamine administration motivated rats to choose a cognitively more demanding option for a higher reward (Cocker, Hosking, Benoit, & Winstanley, 2012; but Hosking, Floresco, & Winstanley, 2015). In keeping with the proposal that dopamine is implicated in the strategic recruitment and/or value-based selection of cognitive control (Boureau, Sokol-Hessner, & Daw, 2015; Hazy et al., 2007), effects of cognitive demand on avoidance learning were shown to depend on striatal dopamine (Cavanagh et al., 2017; Cavanagh, Masters, Bath, & Frank, 2014). More specifically, the presence of response conflict in a Simon task modified learning about action values, such that the value of received rewards was downgraded due to response conflict and a lack of reward after response conflict increased avoidance (Cavanagh et al., 2017, 2014). These effects varied with conditions and manipulations associated with changes in striatal dopamine. For example, they varied as a function of a genetic polymorphism implicating striatal dopamine (DARPR-32), were modulated by a selective D2 receptor agonist (cabergoline) challenge and were altered in patients with Parkinson’s disease, characterized by striatal dopamine depletion (Cavanagh et al., 2017, 2014). A separate line of evidence comes from incentive motivational work, showing that incentive effects on cognitive control vary as a function of striatal dopamine levels in patients with Parkinson’s disease, and healthy volunteers (Aarts et al., 2012, 2014; Manohar et al., 2015). Together, these prior findings raise the question of whether effects of catecholamine manipulations on cognitive control tasks might reflect (in part) changes in value-based learning/choice about cognitive control, in addition to reflecting changes in the ability to execute cognitive control per se.

### The present experiment

In the present experiment, we administered a low, oral dose of MPH to a large group of young healthy volunteers to address our primary question of interest: Does manipulation of catecholamine transmission alter the avoidance of cognitive demand, here task switching? Second, we also investigated effects of MPH on the execution of task switching (performance). To expose individual variation, we obtained indirect estimates of baseline dopamine transmission using behavioural proxy measures of trait impulsivity, associated with dopamine (auto)receptor availability and striatal dopamine release (Buckholtz et al., 2010), as well as working memory span, associated with striatal dopamine synthesis capacity (Cools et al., 2008; Landau et al., 2009). Given prior evidence for greater MPH-induced improvement of performance and learning in higher impulsive participants (Clatworthy et al., 2009; see above), we anticipated greater MPH-induced increases in (the learning of) demand avoidance in higher impulsive participants. Conversely, our hypothesis with regard to working memory capacity was bidirectional, given prior reports of positive, but also negative associations between working memory capacity and cognitive effects of MPH (Mehta et al., 2000; van der Schaaf et al., 2013).

## METHODS

### 2.1 Participants

106 healthy, young adults participated in this study and were recruited via flyers around the campus and the digital participant pool of the Radboud University, Nijmegen. All participants were native Dutch speakers and provided written informed consent to participate in the study. Participants were screened extensively according to pre-defined exclusion criteria (**Supplemental Material 1**).

Data from five participants were incomplete due to medical (irregular heart rate: n = 1, elevated heart rate and nausea: n = 1), and technical (n = 1) problems and drop-outs (n = 2). One additional participant was discarded due to a lack of task understanding (explicitly reported and evidenced by below-chance performance). Thus, the analyses include 100 adult participants (50 women, mean age 21.5, SD = 2.31, range 18 - 28). Two participants had trouble swallowing the capsule such that for one participant the capsule dissolved orally before swallowing and for the other participants content of the capsule was dissolved in water. We assessed whether relevant results were changed when excluding these participants.

We performed a power analysis using G*Power 3.1.9.2 software (Faul, Erdfelder, Buchner, & Lang, 2009; Faul, Erdfelder, Lang, & Buchner, 2007). Previous work from our group had revealed a correlation of 0.74 between a proxy measure of dopamine transmission, working memory capacity, and effects of MPH on reward-learning with 19 participants (van der Schaaf et al., 2013). To be conservative, given the small sample size of that previous study and given that we are using a different experimental task, we anticipated that the true effect size for the present study would be maximally half this size (r = 0.37). Our sample size of 100 (and subsequent subsample of 74: see §3.1) provides 97.6% (92.2%) power to detect such an effect size, for a two-sided test with an alpha-level of 0.05.

Additional demographic and questionnaire information of included participants is reported in Table 1. All procedures were in accordance with the local ethical guidelines approved by the local ethics committee (CMO protocol NL47166.091.13) and in line with the Helsinki Declaration of 1975. The study was also registered with the Dutch National Trial register (trialregister.nl, number NTR4653).

**Table 1.**
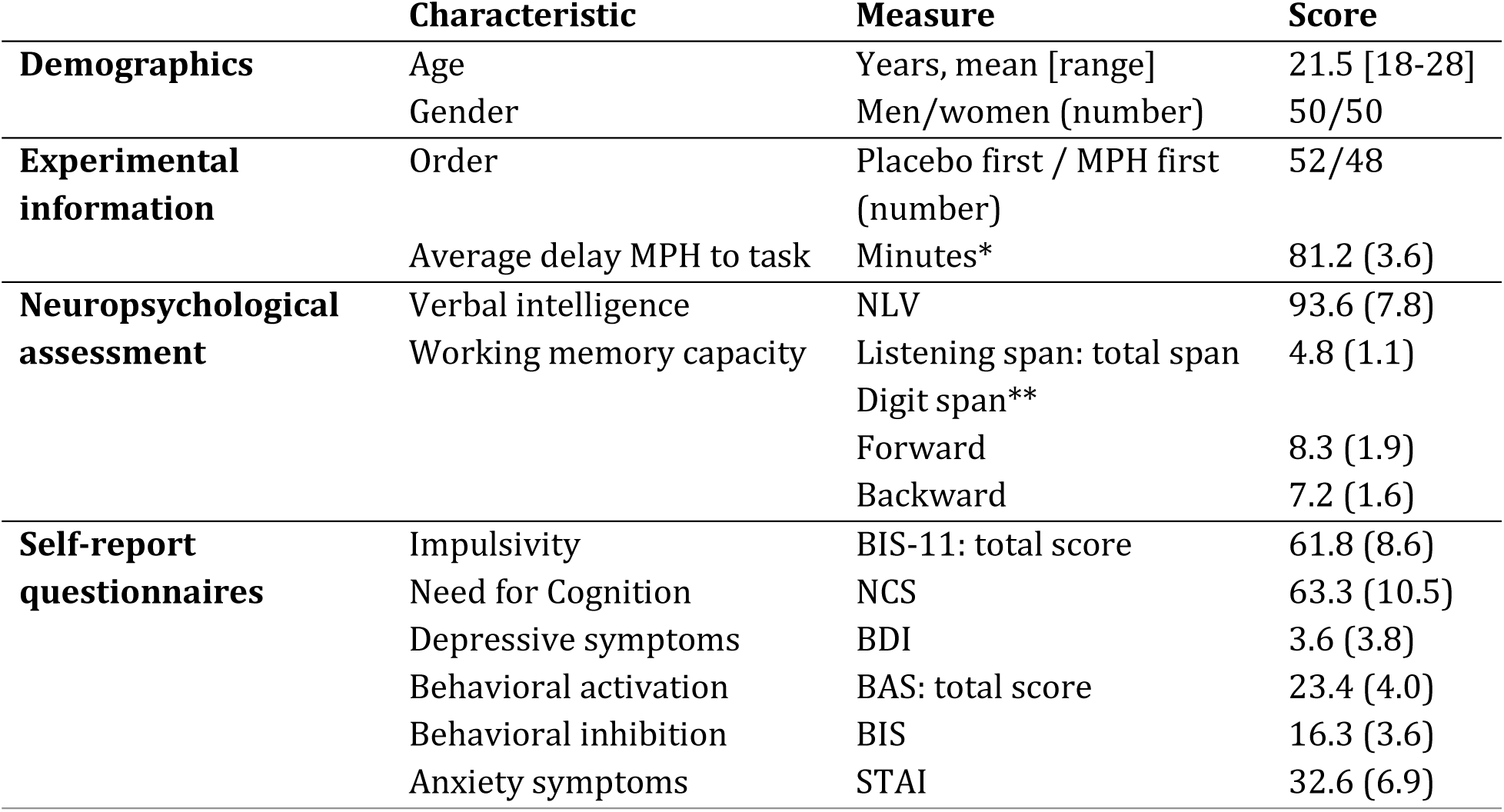
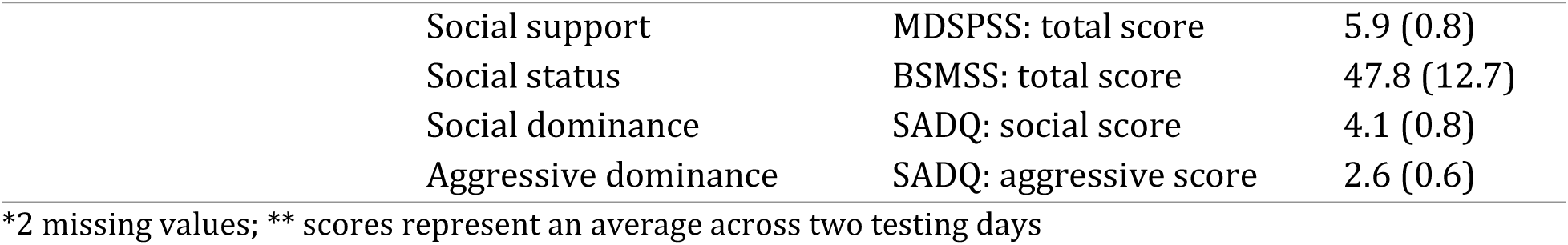
Demographic and background characteristics of participants included in the analysis (n = 100). Questionnaires included the Beck Depression Inventory (BDI; Beck, Steer, Ball, & Ranieri, 1996), Behavioral Inhibition Scale/Behavioral Activation Scale (BIS/BAS; Carver & White, 1994), Spielberger Trait Anxiety Inventory (STAI; Spielberger, Gorsuch, Lushene, Vagg, & Jacobs, 1983), Multidimensional Scale of Perceived Social Support (MDSPSS; Zimet, Dahlem, Zimet, & Farley, 1988), Social and Aggressive Dominance Questionnaire (SADQ; Kalma, Visser, & Peeters, 1993) and Barratt Simplified Measure of Social Status (BSMSS; Barratt, 2006). If not indicated differently, scores represent group averages and the standard deviations between brackets. Reported scores are comparable with observations in healthy populations in earlier reports. Listening span, e.g. Salthouse & Babcock, 1991; Digit span, e.g. van der Schaaf et al., 2014: FW mean = 8.5; BW mean = 7.9; BIS-II, e.g. Buckholtz et al., 2010: mean = 59.5, NCS, e.g. Westbrook et al., 2013: mean = 64.5; BDI, e.g. Schulte-Van Maaren et al., 2013: mean = 3.7; BIS/BAS, e.g. Franken, Muris, & Rassin, 2005: mean BIS = 13.8, mean BAS = 24.5; STAI, e.g. De Weerd et al., 2001: mean ≈ 34; MDSPSS, e.g. Canty-Mitchell & Zimet, 2000: mean = 5.5; BSMSS, e.g. Cook et al., 2014: mean = 49.0, 42.6; SADQ, e.g. Cook et al., 2014: social mean = 4.0, 3.9; aggressive mean = 2.9, 2.7. The verbal IQ estimate (NLV) seems low in this sample (relative to e.g. van der Schaaf et al., 2014: mean = 101). However, we tested a student population and we expect this value to be low due to the outdated character of the test (1991), not accomodating the changes in language use.

### 2.2 Study sessions and pharmacological intervention

A within-subjects, placebo-controlled, double-blind, cross-over design was employed. Participants visited the institute twice for study sessions of around 4.5 hours. The sessions started approximately at the same time of the day across sessions (maximal deviation: 45 minutes), with an interval of one week to 2 months between testing days. After signing an informed consent form, session 1 started with a medical screening (~20 minutes) to check for exclusion criteria (**Supplemental Material 1**). We administered a digit span test (forward and backward; Wechsler, Coalson, & Raiford, 2008), Dutch reading test (NLV; Schmand, Bakker, Saan, & Louman, 1991) and participants received a single oral dose of methylphenidate (MPH; Ritalin^®^, Novartis, 20 mg) on one and a placebo substance on the other day. The order of administration was counterbalanced and double-blind. MPH is known to block transporters of both dopamine (DAT) and noradrenaline (NET), thereby preventing reuptake of catecholamines (Volkow et al., 2002). Plasma concentrations peak after 2 hours with a half-life of 2-3 hours (Kimko, Cross, & Abernethy, 1999). To test participants at maximal plasma levels, participants underwent a cognitive test battery starting 50 minutes after drug intake, including the demand selection task (described in §2.3), the paradigm of primary interest for our research question. The delay between the administration of MPH or placebo and the start of the demand selection task was on average 80.9 (SD = 3.7) minutes. The second testing day was identical to the first one, except that participants performed a listening span test instead of the medical screening (also ~20 minutes, see §2.4). The cognitive test battery consisted in total of six paradigms (Figure 1A). The order of paradigms was constant across sessions and participants, such that a Pavlovian-to-instrumental transfer task (cf. Geurts, Huys, den Ouden, & Cools, 2013) and a social learning task (cf. Cook, Den Ouden, Heyes, & Cools, 2014) preceded the demand selection task on both days, and was followed after a break, by a valenced Go/NoGo learning task (Swart et al., 2017), working-memory task (cf. Fallon et al., 2016), and a probabilistic reversal learning task (cf. den Ouden et al., 2013).

**Figure 1.**
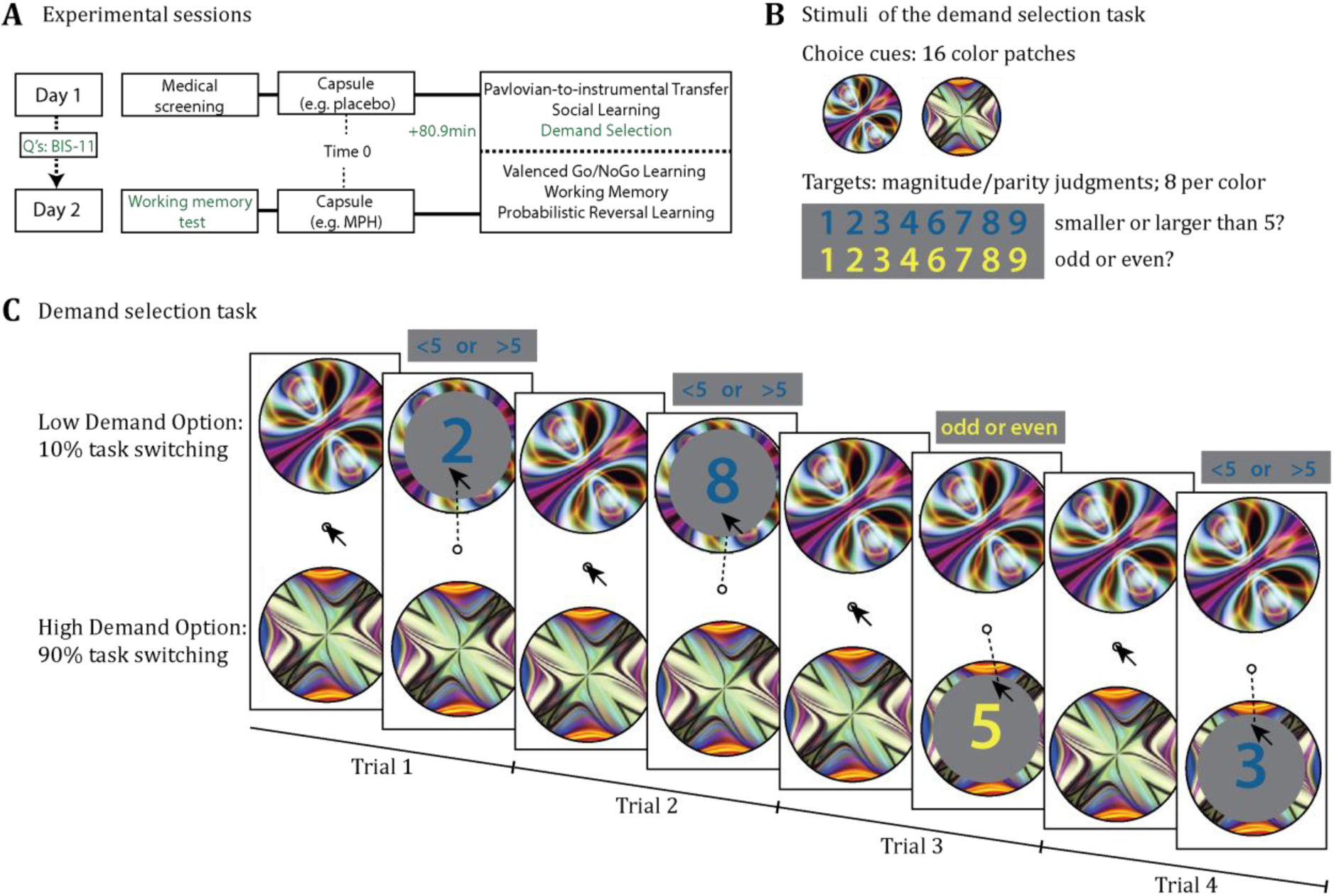
**A** Schematic representation of testing days. Medical screening took place on the first day, a working memory test (i.e. listening span) on the second day. The remaining of the testing days were identical for both days, with methylphenidate (MPH) administration on one day and placebo on the other. Drug administration was followed by a task battery. Between the testing days, participants completed a series of self-report questionnaires, including the BIS-11 impulsiveness inventory. **B** Example stimuli of the demand selection task are presented. Circular colour patches are used as choice cues; the colour of the digits indicates which task had to be executed (magnitude versus parity judgment). **C** Example trial sequence of demand selection task. Participants were shown two colour patches as choice cues. On every trial, participants chose between the two patches, by moving the mouse cursor towards one cue. A digit from 1 to 10 (but not 5) appeared at the target location (putative mouse path indicated here by dashed line). Depending on the colour of the digit, participants either indicated whether the digit was odd or even (i.e. parity judgment for yellow digits), or whether the digit was smaller or larger than 5 (i.e. magnitude judgment for blue digits) by clicking the left or right mouse-button. Responses were self-paced.

For safety reasons blood pressure and heart rate were measured three times throughout the days (start of testing day, before task battery, after task battery). At the same time points, participants’ mood and medical symptoms were assessed using the Positive and Negative Affect Scale (PANAS; Watson, Clark, & Tellegen, 1988), the Bond and Lader Visual Analogue Scales (calmness, contentedness, alertness; Bond & Lader, 1974) and a medical Visual Analogue Scale (symptoms, such as headache or muscle pain; **Supplemental Material 2**). Between the two testing days, participants completed self-report questionnaires at home (see §2.5).

### 2.3 Demand selection task

To assess avoidance of cognitive control, we employed the demand selection task developed by Kool et al. (2010), programmed using the Psychophysics toolbox (Brainard, 1997; Pelli, 1997) in Matlab. Stimuli were 16 random colour fractals used as choice cues and coloured (yellow or blue) digits ranging from 1 to 10 (excluding 5) (Figure 1B). Stimuli were presented on a gray background and responses were made using a pc mouse.

An example trial sequence is presented in Figure 1C. Participants were shown two colour patches as choice cues. After choosing between the two patches, by moving the mouse cursor onto one cue, a digit from 1 to 10 (but not 5) appeared at the center of the chosen cue. Depending on the colour of the digit, the task of the participants was to either indicate whether the digit is odd or even (i.e. parity judgment for yellow digits), or whether the digit is smaller or larger than 5 (i.e. magnitude judgment for blue digits). Judgment was made by clicking the left or right mouse button. After the response, the cursor returned to the center of the screen and the next two choice cues were presented.

Task demand was manipulated by assigning different task switching probabilities to the two choice cues. When choosing one choice cue, the digits switched colours (i.e. task) with respect to the previous trial on 90% of trials. When choosing the other cue, the task repeated on 90% of trials. The option with higher task switching probability represents the more demanding option, based on evidence of task switching requiring extensive cognitive control (Monsell, 2003) and reports of lower accuracy in earlier studies using this task (Kool et al., 2010). The task switching manipulation was unbeknown to the participants. Choice behaviour (i.e. demand avoidance) and performance on the task switching task (i.e. reaction time and accuracy) were the dependent variables of interest.

Participants first practiced 40 trials of only magnitude/parity judgments using the blue and yellow digits as stimuli. Participants were then instructed on the choice task emphasizing that they will choose between two cues repeatedly and the blue or yellow digits will appear at the cue location after they have moved the cursor towards the cue. They were instructed that they could switch between the cues at any point and when they develop a preference for one choice cue, it is fine to keep choosing the same. Instructions were followed by 4 practice choice trials to illustrate the paradigm, but using different cue patches to the actual task. The task consisted of 600 trials, divided across 8 blocks of 75 trials each. Choices and magnitude/parity judgments were not time restricted, i.e. responses were self-paced. The visual identity and location of the 2 choice cues were constant within a block, whereas every new block introduced new choice cues, located in different positions on the screen. The two choice cues were always separated by 180 degrees on an imaginary circle (radius ≈ 11.5 mm) around the center of the screen. The change in visual identity and location of choice cues aimed to prevent motor, location or aesthetic cue preferences confounding the effect of interest (see also Kool et al., 2010). We assessed participants’ awareness of the task switching manipulation using a debriefing questionnaire on the second testing day after task completion. Specifically, we evaluated participants to be aware of the manipulation when they responded positive to the question whether they felt that numbers below one of the two pictures had a tendency to switch between colours more often while the other picture tended to repeat the same colour.

### 2.4 Listening span task

The listening span test (Daneman & Carpenter, 1980; Salthouse & Babcock, 1991) was administered at the beginning of the second test session to obtain an estimate of participants’ working memory capacity, as a putative proxy of baseline dopamine synthesis capacity. During this test, participants listened to pre-recorded sentences and were given two tasks: They answered simple written multiple-choice questions about the content while remembering the last word of each sentence for later recall. The number of sentences on each trial (i.e. the span) increased up to 7 over the course of the task. Three series of the same span were conducted. The trial was coded as successful if the answers to the multiple choice questions were correct and if all last words were remembered and reported in the correct order. Based on participants’ performance a listening span was calculated ranging from 0 to a maximum of 7. The highest level for which two out of the three series were correctly remembered comprised the *basic span*. Half a point was added if one serie of the following span was correctly completed, resulting in the measure of *total span. Total span* and total number of words recalled have both been shown to correlate positively with dopamine synthesis capacity (Cools et al., 2008; Landau et al., 2009) and to predict dopaminergic drug effects (Kimberg et al., 1997; Frank et al., 2006; Cools & D’Esposito, 2011; Kimberg & D’Esposito, 2003; Van der Schaaf et al., 2014).

### 2.5 Questionnaires

A series of questionnaires was completed by participants at home between the two test sessions. The trait impulsivity questionnaire was key to our research question and will be described in more detail below. The other questionnaire data were acquired for exploratory purposes, not pursued here, and are presented in Table 1.

#### Trait impulsivity

The Barratt Impulsiveness Scale (BIS-11; Patton, Stanford, & Barratt, 1995) was administered to assess participants’ degree of trait impulsivity. The scale is a self-report questionnaire, consisting of 30 statements that participants rate on a 4-point Likert scale (“never” to “almost always”). Examples are “I buy things on impulse” or “I am future oriented”. Scores on this questionnaire can range from 30 to 120. The total Barratt score has been found to be associated with reduced dopamine D2/D3 receptor availability in the midbrain, and enhanced dopamine release in the striatum (Buckholtz et al., 2010; Lee et al., 2009) and has been shown to predict effects of MPH on learning (Clatworthy et al., 2009). This measure serves as a second putative proxy of baseline dopamine function for predicting effects of MPH.

### 2.6 Statistical analyses

The experiment was set up to assess effects of MPH on, first, demand avoidance (cue choice) and, second, the execution of task switching (performance). We assessed demand avoidance by analyzing the proportion of participants’ choices of the low demand cue (requiring 10% task switching) versus high demand cue (requiring 90% task switching). Execution of task switching was assessed by analyzing demand costs, which were calculated by subtracting performance (accuracy and (log-transformed) response times (RTs)) on trials on which participants chose the low-versus high-demand option. Following our primary questions, we assessed the effects of MPH on these measures as a function of two putative proxy measures of baseline dopamine function: trait impulsivity, measured with the Barratt Impulsiveness Scale, and working memory capacity, measured with the listening span test. The data were analyzed with mixed-level models using the lme4 package in R (Bates, MÄchler, Bolker, & Walker, 2015; R Core Team, 2013). This allowed us to account for within-subject variability in addition to between-subject variability. Factors drug [MPH vs. placebo] and demand [low vs. high] (for performance only) were within-subject factors, and impulsivity and listening span scores were between-subject factors. Models included all main effects and interactions, except for the interaction between impulsivity and listening span, as our question did not concern this interaction. All models contained a full random effects structure (Barr, 2013; Barr, Levy, Scheepers, & Tily, 2013). P-values reported in the manuscript that pertain to the regressions were estimated using the “esticon” procedure in the “doBy” package which relies on the chi-square distribution (Hojsgaard, 2006). Effects were considered statistically significant if the p-value was smaller than 0.05. We report R^2^ for all models using the “r.squaredGLMM” procedure in the “MuMIn” package to provide a more intuitive estimate. However, note that there is no broad agreement yet about the most appropriate way of R^2^ estimation for mixed-effects models. An overview of the basic models (cue choice, accuracy, RTs) is presented in **Supplemental Table 1**.

#### Response stickiness

Surprisingly, participants displayed extremely high rates of response stickiness, as indexed by low proportions of switching between the cues (note that within blocks all cues had fixed locations). To assess whether the observed choice effects were explained or masked by modulation of response stickiness, we constructed a second and third choice model, which extended the basic model with a stay-regressor and then adding its interaction with MPH. The stay-regressor quantified the degree to which participants’ choices were the same as their choice on the previous trial and allowed us to investigate whether reported drug effects of interest are significant when accounting for (drug effects on) response stickiness. We conducted model comparisons to assess whether the models including response stickiness effects improved our explanation of the data relative to the basic choice model. Model comparison was conducted using the anova function in R, which assesses whether the reduction in the residual sum of squares is statistically significant compared with the simpler model. Results of the winning model will be presented.

To confirm that the MPH-effects on demand avoidance, i.e. our primary choice effect of interest, did not reflect MPH-effects on response stickiness, we also checked whether MPH-effects on response stickiness correlated with MPH-effects on demand avoidance using Spearman correlations (given that the proportion of staying with the same cue violated assumptions of normality and contained outliers) in SPSS 21 (IBM Corp., Armonk, N.Y., USA).

#### Relationship between demand avoidance and task performance

To assess whether MPH-effects on demand avoidance relate to MPH-effects on task performance (accuracy or RTs), we calculated Spearman correlations between the proportion of low demand choices and demand costs (for RT and accuracy) and the MPH-effect on these measures using SPSS 21 (IBM Corp.). To quantify evidence for an absence of effects, we also calculated Bayesian correlations between MPH-effects on these various variables using JASP software (Version 0.7.5; JASP Team, 2016).

## RESULTS

### 3.1 Methylphenidate alters the avoidance of task switching

Cognitive demand was operationalized by two choice options with opposing task switching probabilities (10% vs. 90%). As expected, participants were overall demand avoidant; participants chose the cue with low task switching probability more often than the cue with the high probability (*M* = 0.56, *SD* = 0.13) (Intercept: *X*^*2*^*(1) = 20.70, p < 0.001)*. Demand avoidance was evident both during the placebo and MPH session (Figure 2). A minority of participants (26%) reported during debriefing that they were aware of the fact that one choice cue resulted in more task switches than the other cue.

**Figure 2.**
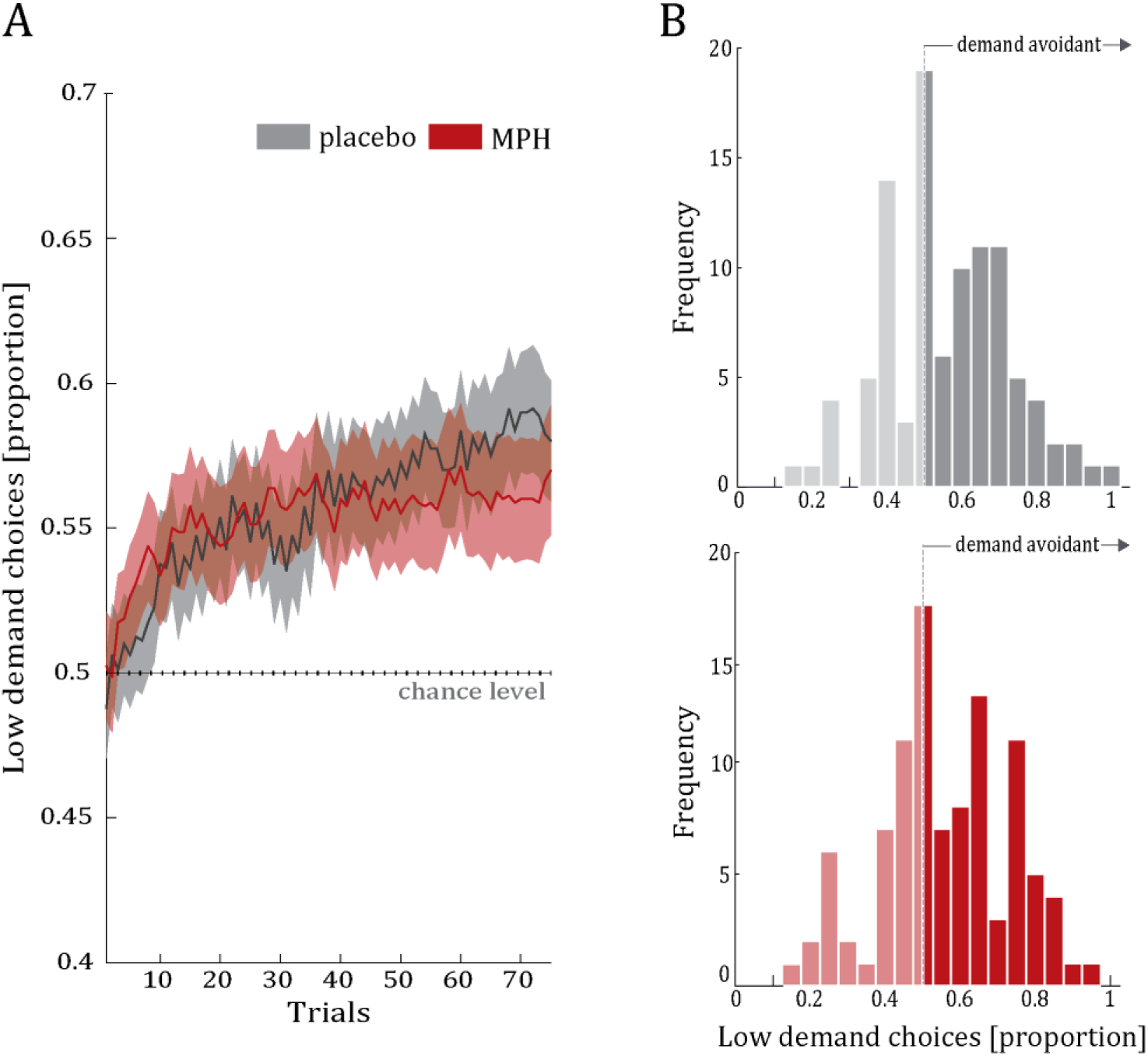
**A** Average proportion of low demand choices as a function of trial, averaged per participant over 8 blocks, for placebo (grey) and methylphenidate (MPH; red) sessions. Data lines represent the group average and shaded area represents standard error of the mean. **B** Histograms of low demand choices in the placebo (top) and MPH (bottom) sessions reveal large individual variability in terms of demand avoidance. Frequency represents number of participants.

Surprisingly, participants exhibited extremely high rates of resonse stickiness, as indexed by the low number of trials on which participants switched between cues (across participants and sessions: *M* = 5.9%, SD = 17.5%) (**Supplemental Figure 1**). Five participants never switched cues in both test sessions. An additional 17 participants never switched cues on one testing day (a further 2 participants switched on every trial). This unexpected high rate of response stickiness in combination with earlier reports of dopaminergic medication effects on response stickiness (Rutledge et al., 2009), led us to ask whether our primary effect of interest on demand avoidance might reflect or be masked by effects on response stickiness. To assess this, we included a stay regressor in the basic choice model (**Supplemental Table 1**). Model comparison with the original basic model lacking the stay regressor showed that a model including a stay regressor (BIC = 54711, marginal R2GLMM = 0.122) explained significantly more variance in choice behaviour than did the basic model (BIC = 150826, marginal R^2^_GLMM_ = 0.004; *X2(1) = 96127, p < 0.001).* However, the model including both, a stay regressor and a regressor for MPH-effect on staying (BIC = 26607, marginal R^2^_GLMM_ = 0.639) explained even more variance than the model without the interaction term (*X ^2^(8) = 28198, p < 0.001*). Therefore we report the results of this extended model below.

Results of this winning model reveal that overall, demand avoidance did not differ between drug sessions (Main effect of Drug: *X*^*2*^*(1) < 0.01, p = 0.964*). However, we hypothesized that effects of MPH on demand avoidance would crucially depend on putative proxies of dopamine transmission, namely trait impulsivity (indexed by total Barratt Impulsiveness Scale score), and/or working memory capacity (indexed by total listening span). As predicted, MPH-effects on demand avoidance varied significantly as a function of trait impulsivity (Drug x Impulsivity: *X*^*2*^*(1) = 5.33, p = 0.021)*. The direction of this effect was positive with greater MPH-induced increases in demand avoidance in more impulsive participants (Figure 3). The interaction between working memory capacity and the effect of MPH on demand avoidance was only trending towards significance (Drug x Listening span: *X*^*2*^*(1) = 2.91, p = 0.088*). We therefore focus further analyses on trait impulsivity, while reporting further analyses as a function of working memory capacity in the **Supplemental Results 1**.

**Figure 3.**
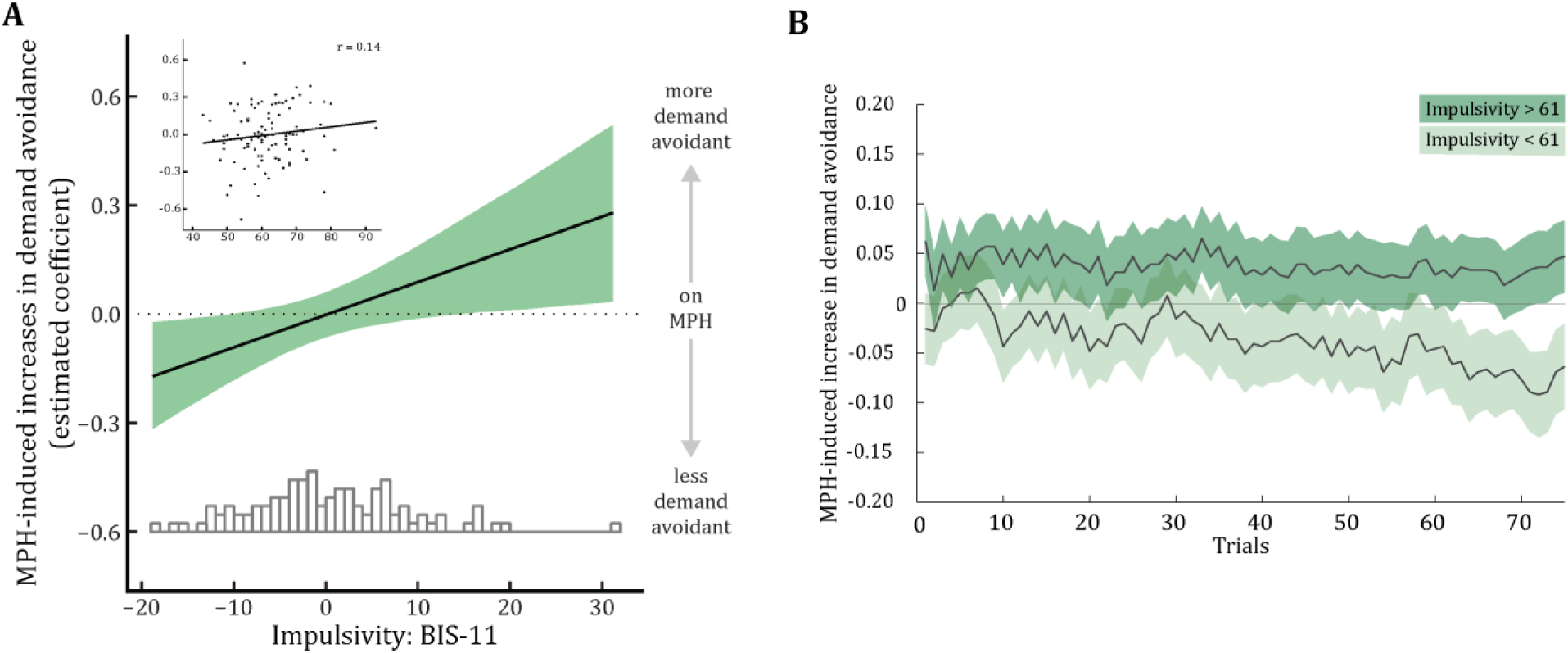
Methylphenidate-effect on demand avoidance as a function of participants’ trait impulsivity (BIS-11) scores. **A** Line represents model-based estimated coefficients of MPH-effect on demand avoidance as a function of (z-scored) trait impulsivity scores. Shaded area represents simulated 95% confidential intervals of the coefficients. The inset shows the raw data: drug effect for every participant (n = 100) across trials as the difference in the proportion of low demand choices (MPH - placebo) as a function of trait impulsivity. **B** Trial-by-trial drug effect averaged across 8 blocks, and across participants (n = 100) of low (n =49) versus high (n = 48) trait impulsivity groups as a function of trial. 3 participants with scores equal to the median are not included. Shaded areas represent standard error of the difference. See **Supplemental Figure 3** for the impulsivity-dependent effect of MPH as a function of trial number for placebo and MPH separately.

In addition, results of the winning model reveal, apart from a main effect of staying with the previously chosen option (staying: *X*^*2*^*(1) = 291.16, p < 0.001*), that MPH also affected staying (Drug x Stay: *X*^*2*^*(1) = 7.65, p = 0.006).* MPH increased response stickiness relative to placebo. Complete statistics of this choice model are presented in **Supplemental Table 2**.

To confirm that these effects of MPH on response stickiness could not explain the impulsivity-dependent demand avoidance effects, we also investigated whether there was any correlation between MPH-effects on the proportion of staying with the same choice cue and MPH-effects on demand avoidance. There was no such correlation (low demand choices mph – pla & proportion staying mph – pla: *rs* = 0.12, *p* = 0.240;), with Bayesian correlation analysis showing substantial evidence for the null effect (BF_10_= 5.14).

Finally, the size of our sample allowed us to assess whether the impulsivity-dependent effects remained present when excluding participants who appeared to use explicit choice strategies, i.e. failed to explore the choice options at all, either in one (n = 17) or both sessions (*n* = 5), and those who switched between choice cues on every trial, either in one (n = 1) or both sessions (n = 1). We also excluded those participants for whom the capsule dissolved (orally or in water) before swallowing (n = 2, one of those was also a sticky participant) as well as one participant whose score on the BIS-11 deviated more than 3 standard deviations from the mean. Analysis of this smaller dataset (*n* = 74) confirmed the effects obtained from the analysis of the larger sample: MPH altered demand avoidance significantly as a function of trait impulsivity (Drug x Impulsivity: *X*^*2*^*(1) =* 5.80*, p =* 0.016; **Supplemental Figure 2; Supplemental Table 3**).

In sum, above control analyses show that observed MPH-effects on demand avoidance are robust, also when taking into account MPH-effects on response stickiness or excluding problematic participants. Furthermore, a correlation analysis suggests that MPH-effects on response stickiness and demand avoidance are independent.

### 3.2 Avoidance of task switching does not reflect poor performance

Following every cue choice (10% vs. 90% task switching probability), participants were presented with a parity/magnitude judgment task. Overall accuracy was high in this number judgment task (*M* = 0.97, *SD* = 0.04) and, as expected, participants were sensitive to the task switching manipulation. They performed better when the task repeated with respect to the previous trial than when they were presented with a task switch, evidenced by higher accuracy (*M* = 0.01, *SD* = 0.02) (*X*^*2*^*(1)* = 20.43, *p* < 0.001) and faster RTs (*M* = 0.36, *SD* = 0.29) (*X*^*2*^*(1)* = 119.70, *p* < 0.001) (Table 2). The improved performance for task repetitions consequently affected performance for the two cue options: Participants performed better on trials on which they chose the low demand (10% task switching) relative to high demand (90% task switching) cue (accuracy demand cost: *X*^*2*^*(1)* = 20.93, *p* < 0.001; RT demand cost: *X*^*2*^*(1)* = 535.73, *p* < 0.001).

**Table 2.** Mean values (and standard deviations) of choice proportions and performance on the magnitude/parity judgment task (i.e. accuracy, response times, switch and demand costs) for placebo and methylphenidate sessions.

| **Task component** | **Variable** | **Placebo** | **MPH** |
| --- | --- | --- | --- |
| **Demand avoidance**<br>**(i.e. choices)** | Low Demand choice<br>(proportion) | 0.56 (0.17) | 0.55 (0.18) |
|  | High Demand choice<br>(proportion) | 0.44 (0.17) | 0.45 (0.18) |
| **Performance** | Accuracy<br>(in proportion correct) | 0.97 (0.03) | 0.97 (0.07) |
|  | Task repeat | 0.97 (0.03) | 0.97 (0.06) |
|  | Task switch | 0.96 (0.04) | 0.96 (0.07) |
|  | Switch cost | 0.01 (0.02) | 0.01 (0.02) |
|  | Low demand | 0.97 (0.03) | 0.97 (0.06) |
|  | High demand | 0.96 (0.04) | 0.96 (0.07) |
|  | Demand cost | 0.01 (0.02) | 0.01 (0.02) |
|  | Response times<br>(in sec) | 1.06 (0.33) | 0.99 (0.26) |
|  | Task repeat | 0.89 (0.24) | 0.84 (0.21) |
|  | Task switch | 1.25 (0.44) | 1.16 (0.33) |
|  | Switch cost | 0.36 (0.29) | 0.32 (0.19) |
|  | Low demand | 0.90 (0.24) | 0.86 (0.21) |
|  | High demand | 1.25 (0.44) | 1.16 (0.33) |
|  | Demand cost | 0.35 (0.28) | 0.30 (0.20) |

There were no effects of MPH, relative to placebo, on the size of these switch or demand costs, when assessed across the group as a whole (Drug x Demand for RTs: *X*^*2*^*(1)* = 0.75, *p* = 0.387, for accuracy: *X*^*2*^*(1)* = 1.20, *p* = 0.274; Drug x Switch for RTs: *X*^*2*^*(1)* = 0.61, *p* = 0.434, for accuracy: *X*^*2*^*(1)* = 1.91, *p* = 0.167). In contrast to the altered demand avoidance, the effect of MPH on the demand cost did not vary as a function of trait impulsivity (for RTs:Drug x Impulsivity x Demand: *X*^*2*^*(1)* = 0.29, *p* = 0.590, Figure 4A; for accuracy: Drug x Impulsivity x Demand: *X*^*2*^*(1)* = 0.001, *p* = 0.968, Figure 4B).

**Figure 4.**
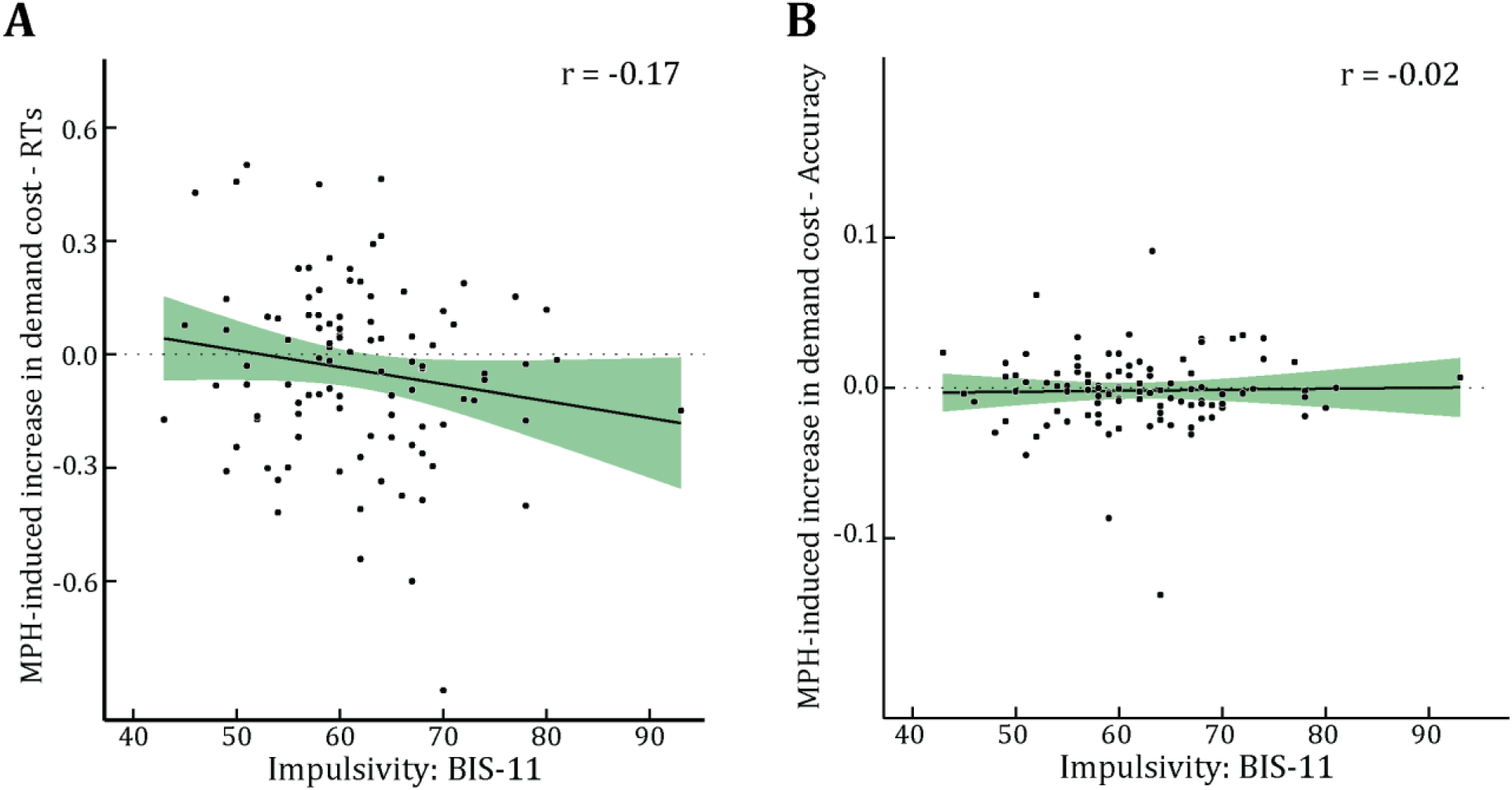
Drug effects on performance costs between high and low demand choices. Data points represent methylphenidate (MPH) effects on average demand cost (MPH - placebo) for each participant (n = 100) for **A** response times (in seconds) as a function of trait impulsivity (BIS-11) and **B** accuracy (in proportion correct) as a function of trait impulsivity (BIS-11). Shaded areas represent standard errors of the mean. Both effects are not statistically significant.

Independent of demand and baseline-measures, MPH increased overall accuracy (Main effect of Drug: *X*^*2*^*(1)* = 8.97, *p* = 0.003), and trended towards speeding up responses (Main effect of Drug: *X*^*2*^*(1)* = 2.98, *p* = 0.084). Interestingly, these MPH-induced response time (but not accuracy) changes did depend on trait impulsivity (Drug x Impulsivity: *X*^*2*^*(1)* = 7.28, *p* = 0.007), with greater MPH-induced decreases in response times in more impulsive participants. Complete statistics of the basic performance models are presented in **Supplemental Table 4**.

This pattern of findings suggests that MPH-induced demand avoidance cannot be explained by reduced performance under MPH (i.e. avoidance of failure). MPH increased demand avoidance in more impulsive participants despite MPH-induced speeding of responding and unaffected accuracy (Drug x Impulsivity: *X*^*2*^*(1) < 0.01, p =* 0.747), also not as a function of demand (Figure 4B).

Although the reported findings above suggest that performance cannot explain the MPH-induced demand avoidance, we further assessed the potential association with a direct correlation. In other words, we tested whether participants who avoided demand more, did so because the task had become more difficult for them. More specifically, we computed correlations between demand costs (accuracy and RT) and demand avoidance. In line with our reasoning above, the MPH-effect on demand costs (demand cost MPH - PLA) did not correlate with the drug effect on demand avoidance (low demand MPH - PLA) and even provided evidence, though weak, for the absence of the correlation (accuracy: *rs* = 0.14, *p* = 0.167, BF_01_= 4.80; RT: *rs* = -0.10, *p* = 0.330, BF_01_ = 2.60).

In sum, analyses of performance data and correlations between performance and demand avoidance provide evidence that observed MPH-effects on demand avoidance are unlikely to be explained by performance changes. This suggests that while the actual performance of the task did not change, this demand was evaluated differently (indexed by degree of demand avoidance). Bayesian analyses provided evidence for independence of the demand avoidance and performance effects.

## DISCUSSION

In this study, we investigated whether prolonging catecholamine transmission alters choices about whether or not to recruit cognitive control (i.e. demand avoidance). Specifically, we hypothesized that challenging the catecholamine system would alter the avoidance of cognitive demand. We tested this hypothesis by assessing the effects of acute administration of oral MPH (20mg), a potent blocker of catecholamine transporters, on task switching avoidance using a demand selection task (Kool et al., 2010). A large sample of young healthy participants (n = 100) was tested to expose well-established individual differences in the response to such catecholaminergic drugs (Cools & D’Esposito, 2011). Given the well-established observation that drug effects vary across individuals as a function of baseline levels of dopamine, we obtained indices of trait impulsivity and working memory capacity, both previously associated with dopamine transmission (Cools et al. 2008; Landau et al., 2009; Buckholtz et al., 2010; Dalley et al., 2007; Lee et al., 2009; Kim et al., 2013; Reeves et al., 2012). As predicted, MPH altered the avoidance of task switching, without changing the execution of task switching itself. Notably, this effect was isolated when taking into account trait impulsivity.

### General demand avoidance effects

On average and across sessions, participants chose the low demand option more often than the high demand option, which indicates that our paradigm was sensitive to our construct of interest, i.e. demand avoidance, consistent with prior studies using this or very similar paradigms (Kool et al., 2010; Kool, McGuire, Wang, & Botvinick, 2013; McGuire & Botvinick, 2010). Moreover, as previous, demand avoidance was observed despite most participants reporting not to be aware of the demand manipulation. Thus we replicate previous observations that anticipated cognitive demand contributes to decision-making, so that decisions are made, partly, in order to minimize demands for exertion or work, a principle sometimes referred to as the law of less work (see also Botvinick, 2007; Westbrook et al., 2013). However, the average proportion of low demand choices was somewhat lower in our study compared with previous work (e.g. Kool et al 2010; see results section; but Gold et al., 2015). It is possible that this reflects the fact that our participants exhibited very high rates of response stickiness, perhaps due to a relatively reduced engagement with or enhanced avoidance of performing the choice task itself.

### Methylphenidate alters demand avoidance in a baseline-dependent manner

Our key finding was that MPH affects demand avoidance, but that these effects varied as a function of trait impulsivity, with greater MPH-induced increases in demand avoidance in more, relative to less, impulsive participants. Much progress has been made recently in our understanding of the (psychological, neurochemical, and neural) mechanisms of our motivation to avoid cognitive demand (e.g. Chong et al., 2017; Cools, 2016; Shenhav et al., 2017; Westbrook & Braver, 2016). Here, we focus on the psychological and chemical neuromodulatory mechanisms of demand avoidance.

Most generally, the motivational control of goal-directed behaviour is well established to depend on the learning of the value and cost of our actions (Dickinson & Balleine, 1994). Factors that have been suggested to contribute to the motivational control of specifically our cognitive actions include the learning of time (opportunity) costs (Boureau et al., 2015; Kurzban et al., 2013), of intrinsic effort costs related to conflict (Cavanagh et al.,2014; Kool et al., 2013), of error likelihood or performance failure (Dunn & Risko, submitted) and/or a combination of these factors (Dunn, Lutes, & Risko, 2016; Shenhav et al., 2017).

We began to address the psychological mechanism underlying our effect on demand avoidance by asking whether it can be attributed to indirect effects on performance costs (error or RT). This is unlikely in the current dataset, for the following reasons. First, there was evidence for an absence of correlations between MPH-induced demand avoidance and MPH-induced performance effects (i.e. demand costs in error rates and RTs) across participants. Second, demand costs were not modulated by MPH. Finally, in more-relative to less-impulsive participants, MPH increased demand avoidance, but actually improved task performance in terms of response speed. Thus the MPH-induced changes in demand avoidance are unlikely to reflect indirect effects of modulation of (perceived) performance failure.

Instead, we hypothesize that MPH might alter demand avoidance via modulating an intrinsic, or opportunity cost of effort. This hypothesis concurs generally with recent work showing that the effect of demand, manipulated by response conflict, on reward versus punishment learning varies with pharmacological dopamine receptor stimulation as well as individual genetic variation in dopamine transmission (Cavanagh et al., 2014). It might be noted that the present study was not set up (and, given high response stickiness rates, did not allow us) to disentangle the degree to which the MPH-effect on demand avoidance reflects learning (or choice) based on reward (effort relief) or punishment (effort cost).

MPH prolongs catecholamine transmission in a nonspecific manner by targeting both dopamine and noradrenaline transporters (Kuczenski & Segal, 2001; Scheel-Krüger, 1971). Therefore, a key remaining open question is whether the effects of MPH, reported here, reflect modulation of dopamine or noradrenaline. We hypothesize, in part based on the work by Cavanagh et al. (2014), reviewed above, that our effect of MPH on demand avoidance reflects modulation of striatal dopamine. This concurs with a recent study reporting striatal dopamine increases after administration of a low-dose of MPH (Kodama et al., 2017) and also with our previous finding that the effects of MPH on reward-versus punishment-learning resembled that of the selective dopamine receptor agent sulpiride, which has selective affinity for D2 receptors that are particularly abundant in the striatum (Janssen et al., 2015; van der Schaaf et al., 2014). Moreover, it is generally consistent with prior work, demonstrating a key role for (striatal) dopamine in physical effort-based choice (Buckholtz et al., 2010; Hosking et al., 2015; Salamone et al., 2016; Wardle, Treadway, Mayo, Zald, & de Wit, 2011), although a recent study failed to observe modulation by the selective dopamine antagonists eticlopride and SCH23390 of the willingness to exert cognitive effort (Hosking et al., 2015). Finally, the dopamine hypothesis coincides with our finding that the effect of MPH depended on trait impulsivity, which implicates increased drug-induced dopamine release (Buckholtz et al., 2010) and changes in D2/D3 receptor availability, although there is discrepancy with regard to the direction of the association between trait impulsivity and D2/3 receptor availability (Buckholtz et al., 2010; Dalley et al., 2007; Lee et al., 2009; Kim et al., 2013; Reeves et al., 2012).

Future studies are needed to test the hypothesis that MPH alters demand avoidance via affecting dopamine rather than noradrenaline transmission, for example using a MPH administration design in which participants are pretreated with a selective dopamine receptor antagonist prior to receiving MPH or in which effects of MPH are compared with those of atomoxetine, which leaves unaltered striatal dopamine transmission. This is especially pertinent because of the well-established link between the locus coeruleus– norepinephrine system and mental fatigue (Berridge & Waterhouse, 2003) and the implication of this system in task-related decision processes and optimization of task performance (Aston-Jones & Cohen, 2005). Moreover, recent empirical evidence indicates a key role for noradrenaline in task engagement and meta-cognitive regulatory functions (Hauser et al., 2017; Hopstaken, van der Linden, Bakker, & Kompier, 2015).

### Methylphenidate does not alter the execution of task switching

Unlike MPH-effects on demand avoidance, there were no effects of MPH on the actual performance of the task, as indexed by performance costs in accuracy or response times. Taking into account trait impulsivity did not reveal such an effect of MPH on demand (or switch) costs either. This contrasts with previous work, which showed an amphetamine-induced improvement of task switching (Samanez-Larkin & Buckholtz, 2013). This discrepancy might reflect the fact that the current paradigm was not optimized for measuring rapid task switching. In our paradigm, the number judgement trials were separated by the choice events, thus likely reducing sequential effects like task switching, as subjects needed to switch already between the number judgment task and choices. As a result, the paradigm is likely less sensitive to subtle effects of chemical neuromodulatory effects than were the rapidly paced task switching paradigms used previously (Samanez-Larkin & Buckholtz, 2013).

Across high and low demand trials, MPH speeded responding in high-versus low-impulsive participants, consistent with dopamine’s well-established role in nonspecific behavioural activation and invigoration of responding (Niv et al., 2007; Robbins & Everitt, 2007). Importantly, the overall speeding of responses was not accompanied by an impulsivity-dependent decrease in accuracy, speaking against a shift in the speed-accuracy tradeoff or more sloppy responding and putatively in favour of cognitive enhancement.

In line with various reports on MPH’s potential to enhance cognition after single, low-dose administration (Berridge & Arnsten, 2015; Linssen et al., 2014; Spencer, Devilbiss, & Berridge, 2015), in this study MPH improved overall accuracy of responding on the task switching task, irrespective of demand or baseline measures.

### Response stickiness

We were surprised about the high levels of response stickiness in the choice task and, regardless of its origin, carefully scrutinized our data to assess the possibility that MPH-effects on stickiness reflect or mask our MPH-effect of interest on demand avoidance. For example, an increase in stickiness might have resulted in a failure to explore and to assign high or low effort costs to the two options. This is particularly pertinent, because we observed in the current data that MPH increased response stickiness across participants, and that a logistic regression model which included (MPH-effects on) response stickiness explained more variance than did a model without response stickiness. Moreover, consistent with our effect, prior work has shown that dopaminergic medication in Parkinson’s disease increased response stickiness during a reinforcement learning task (Rutledge et al., 2009; see also Beeler, 2012; Beeler, Faust, Turkson, Ye, & Zhuang, 2016). In fact, it is highly unlikely that the impulsivity-dependent effect of MPH on avoidance reflects modulation of response stickiness. First, the logistic regression model which controlled for response stickiness revealed significant effects of MPH on demand avoidance as a function of impulsivity, even when variability in stickiness was removed. Second, there was substantial evidence for an absence of a correlation between the effect of MPH on demand avoidance and that on response stickiness. Third, supplementary analyses revealed that the same effect remained significant after excluding participants who failed to explore the choice cues. Together, these supplementary control analyses strengthened our confidence in the dependence of the MPH-effect on trait impulsivity, generally consistent with previous results showing greater effects of MPH on learning in high versus low-impulsive participants (Clatworthy et al., 2009).

### Implications

The measure of trait impulsivity was primarily included in this study for its established relation with baseline dopamine transmission. However, impulsivity is also a clinically relevant dimensional trait implicated in multiple psychiatric disorders, such as (drug) addiction or ADHD. One direct implication of our findings is that while MPH may enhance (task-nonspecific) performance in high-impulsive participants (e.g. by altering response speed), consistent with its performance enhancing effect in ADHD, it may also reduce their motivation for (i.e. value-based learning about) cognitive control. This effect on the avoidance of control might seem paradoxical, given that MPH has been shown to i) remedy cognitive control problems in ADHD patients, who are characterized by high levels of impulsivity (Aron et al., 2003; Cepeda et al., 2000; Coghill et al., 2013; Faraone & Buitelaar, 2010; Leonard et al., 2004; Mehta, Goodyer, & Sahakian, 2004b), ii) to improve performance on attention tasks in high-impulsive rats (Puumala et al., 1996; Robbins, 2002) and iii) to enhance task switching in healthy volunteers (Samanez-Larkin & Buckholtz, 2013). However, none of these studies examined the motivation or willingness to recruit or avoid cognitive control. The present results indicate that any cognition and performance enhancing effects of MPH might be accompanied by an (undermining) effect of MPH on the motivation to exert cognitive control.

A second implication of the present findings is that the cognitive control effects of disorders that implicate the catecholamine system, such as ADHD or Parkinson’s disease might (in part) be consequences of changes in the motivation to avoid cognitive control, rather than reflecting changes in the ability to execute control per se (Schneider, 2007). This generally concurs with a characterization of ADHD and Parkinson’s disorder as disorders of the will.

Finally, in line with recent work by Kool and colleagues (2017), our results raise the hypothesis that previously established effects of dopamine on the reliance on cognitively effortful (e.g. model-based versus model-free) behavioural control strategies (Deserno, Huys, Boehme, Buchert, & Heinze, 2015; Wunderlich, Smittenaar, & Dolan, 2012) reflect partly modulation of cost-benefit decision-making rather than ability to execute such strategies.

### Conclusion

We demonstrate that prolonging catecholamine transmission by MPH administration altered the avoidance of cognitive demand in healthy volunteers. These effects were isolated by taking into account individual differences in trait impulsivity. Control analyses support our conclusion that reported MPH-effects on demand avoidance are likely results of a modulation of value-based decision-making and not an indirect consequence of modulation of task performance.

### Context

The study setup is based on earlier work in our group and the field as a whole. It builds on two key-findings: we have established that (1) catecholaminergic drugs can alter performance in various domains, such as working memory (Fallon et al., 2016), reversal learning (van der Schaaf et al., 2013) and selective attention (Ter Huurne et al., 2015) and that (2) individudal differences of such intervention effects likely reflect variability in baseline catecholamine functioning (Cools & D’Esposito, 2011).

In parallel, recent theoretical and modeling work has advanced our insights in the field of motivated cognition and therefore informed our research questions. Specifically, it has been proposed that cognitive control recruitment is not purely limited by our ability to execute cognitive control, but is also a function of value-based learning and decision-making, similar to a willingness to invest principle (e.g. Cools, 2016; Shenhav et al., 2017; Westbrook & Braver, 2016), which was hypothesized to be sensitive to changes in the catecholamine system. The aim of this study was therefore to test for the first time in humans whether cognitive control recruitment, or the avoidance thereof, was altered by a catecholamine challenge, further emphasizing the relevance of quantifying motivational in addition to performance aspects of cognition.

## Acknowledgement

We thank Dr. Wouter Kool for sharing the task code for the demand selection task, Monique Timmers and and Peter Mulders for medical assistence during data acquisition and Dr. Sean Fallon for advice on study setup and ethical approval procedure.

